# Nanoparticle Shape Governs Immunomodulation of MUC1 Antigen to Develop Anti-cancer Vaccine

**DOI:** 10.1101/2021.09.29.460739

**Authors:** Suraj Toraskar, Preeti Madhukar Chaudhary, Raghavendra Kikkeri

## Abstract

T-cell-dependent immunomodulation of carbohydrate antigens under benign conditions is the most promising approach for carbohydrate-based vaccine development. However, to achieve such adaptive immune responses, well-defined multifunctional nanocarriers loaded with immunogenic materials must be explored. Current efforts to use gold nanoparticles (AuNPs) as antigen carriers in vaccine development have conveniently introduced considerable diversity. Here, we show that the shape of AuNPs markedly influences carbohydrate-based antigen processing in murine dendritic cells (mDCs) and subsequent T-cell activation. In the study, CpG-adjuvant coated sphere-, rod-, and star-shaped AuNPs were conjugated to the tripodal Tn-glycopeptide antigen to study their DC uptake and the activation of T-cells in the DCs/T-cell co-culture assay. Our results showed that sphere- and star-shaped AuNPs displayed relatively weak receptor-mediated uptake but induced a high level of T helper-1 (Th1) biasing immune responses compared with rod-shaped AuNPs, showing that receptor-mediated uptake and cytokine secretion of nanostructures are two independent mechanisms. Significantly, the shapes of AuNPs and antigen/adjuvant conjugation synergistically work together to modulate the effective anti-Tn-glycopeptide immunoglobulin (IgG) antibody response after *in vivo* administration of the AuNPs. These results show that by varying the shape parameter, one can alter the immunomodulation, leading to the development of carbohydrate vaccines.

## Introduction

T-cell-dependent (TD) antibody induction is crucial for successfully designing long-lasting immune responses and vaccine development.^1-5^ In TD activation, B-cells proliferate, produce immunoglobulin class switching (immunoglobulin M [IgM] to immunoglobulin G [IgG]) and generate memory B-cells, resulting in high affinity with longer living antibodies.^6^ Hence, traditional vaccines are composed of antigens, carrier proteins, and co-stimulants, such as toll-like receptor (TLR) adjuvants, to enhance anti-antigen immune responses. Despite sustained efforts by immunologists, there remains a need for a new, versatile method to bring about an effective immune-modulatory system, particularly for such weak immunogenic antigens as carbohydrates.^7-9^

Carbohydrates on the cell surfaces of pathogens and cancer cells are promising antigens for vaccine development.^10, 11^ However, because of the immunodominance of carrier proteins over antigens, carbohydrate antigen-carrier protein conjugation methods have typically failed to induce robust humoral and cellular immune responses against self-antigenic tumor-associated carbohydrate antigens (TACAs).^12,13^ Alternatively, nanostructures, virus-like particles, liposomes, and polymers have been used as antigen-carrier platforms to avoid antibody production on scaffolds.^14-19^

Among the various nanostructures used, gold nanoparticles (AuNPs) have received increasing attention because of their nontoxicity and easily tunable physical properties.^20-23^ Recently, Corzana and coworkers reported synthesizing AuNPs carrying human mucin 1 (MUC1)-like antigens bearing *O/S*-glycosidic linkage-displayed immunogenic activity.^24^ Westerlind and coworkers investigated AuNPs functionalized with chimeric peptides, containing the MUC1-derived glycopeptide sequence and the P30 sequence of T-cell epitope for selective antibody responses.^25^ Similarly, Barchi and coworkers reported developing AuNPs bearing TACAs to develop adjuvant-free immune responses.^26^ Furthermore, Barchi and coworkers reported spherical AuNPs bearing 28-mer MUC4 antigens showing IgG responses.^27^ However, most AuNPs used for glycan immunogenicity are spherical nanostructures carrying full-length glycopeptides. Thus, the influence of AuNPs with different shapes on the immunomodulation of carbohydrate antigens has not yet been examined. Moreover, shape-dependent immune modulation is fundamental to developing new biomaterials.

Recently, viral antigens have been encapsulated on different shapes of AuNPs to alter immune responses and aid in vaccine development.^28, 29^ However, a systematic investigation of nanostructure shapes modulating the immune response of carbohydrate antigens has not been described. Herein, we report on the synthesis of sphere-, rod-, and star-shaped AuNPs bearing an active part of MUC1 antigen glycopeptide and a CpG adjuvant to study the shape-dependent immune modulation of the MUC1 antigen. Extensive imaging and fluorescence-activated cell sorting (FACS) analysis with murine dendritic cell (mDC) uptake and cytokine secretion in mDC/T-cell co-culture assay established the relation between the physical properties of AuNPs in receptor-mediated mDC uptake and T-cell activation. Finally, we assessed *in vivo* humoral immune responses to show shape-dependent anti-MUC1 antibody production and representation of a novel platform that can be used for vaccine development.

## Results and Discussion

### Synthesis of antigen/fluorescent conjugate

With the goal of creating a new nanostructure for glyco-immunomodulation, MUC1, which is overexpressed in cancer cells,^30-32^ and the CpG-ODN adjuvant were selected as the immunogenic substrates. The rationale for choosing CpG-ODN as an adjuvant is that it has a strong agonistic nature for TLR9, which induces Th1-biased immune responses.^33, 34^ Studies have been conducted to target MUC1 using various innovative platforms,^35-45^ findings that the Ala-Pro-Asp-Thr-Arg-Pro (APDTRP) region is the minimum immunogenic epitope of MUC1 responsible for strong tumor cell recognition.^46^ Based on this information, we prepared a Tn-glycopeptide (TnG) with an amine linker for nanostructure-mediated immune activity. We used solid-phase peptide synthesis and **5** as the Tn–amino acid residue to synthesize the glycopeptide (Scheme 1). Compound **5** was synthesized *via* glycosylation of the Fmoc-L-threonine amino acid derivative to **2** using trimethylsilyl trifluoromethanesulfonate (TMSOTf) as a promoter, followed by reducing the azide group of **3** through Zn/AcOH and subsequent acetylation using acetic anhydride in tetrahydrofuran (THF) solvent. Finally, the *t-*butyl ester group of **4** was deprotected in acidic condition to obtain compound **5**. Then, using rink amide resin and Fmoc-chemistry, TnG **6** was synthesized with an amine linker using 2-(1H-benzotriazol-1-yl)-1,1,3,3-tetramethyluronium hexafluorophosphate/hydroxybenzotrizole (HBTU/HOBt) as the coupling agent. The final peptide was purified using high-performance liquid chromatography (HPLC) and characterized with nuclear magnetic resonance (NMR) and mass spectroscopic techniques. The fluorescein-linker (**F-1**) was synthesized using the previous procedure with slight modifications to obtain a dithiol linker (Scheme S3).^47^ We envision conjugating this Tn glycopeptide **6** and florescent tag to well-defined tripod **7** to obtain a multivalent system.

The synthesis of tripod **7** was carried out using a previously reported procedure.^48^ Further, **8** was obtained by *N,N’*-dicyclohexylcarbodiimide (DCC) coupling of 5-(Boc-amino)pentanoic acid with free amine of **7** followed by Boc deprotection and reaction with DL-α-lipoic acid *N*-hydroxy succinimide. Ester hydrolysis of **8** was followed by active ester formation, giving compound **9**. Later one of the active ester group of tripod **9** was selectively replaced by *N*-(2-aminoethyl)maleimide, and the remaining two active esters were replaced by TnG **6**. Next, the Michael addition of the FITC linker to the maleimide group led to the final tripod **10**. The final compound was purified by HPLC and characterized by mass spectroscopy.

### Characterization of antigen/adjuvant-coated nanostructures

To assess the interplay between the Tn antigen, CpG adjuvant, and shapes of AuNPs in immunomodulation, we first synthesized two different sizes of the spheres (**S-1**: 20 nm and **S-2**: 45 nm) and rods (**R-1**: 20 × 6 nm and **R-2**: 46 × 14 nm), and single-size star-shaped gold nanoparticles (**St**: 70 nm spike-to-spike), using the previously reported protocol.^49-53^ We chose the above sizes of AuNPs because it has been proven that particles of 10–100 nm can regulate effective lymph node targeting and immunomodulation.^54^ The NPs’ size and shape were confirmed by transmission electron microscopy (TEM) (Figure 1) and ultraviolet-visible (UV-vis) absorption (Figure S1). The spheres displayed localized surface plasmon resonance (LSPR) peaks at 524 nm and 548 nm, corresponding to two sizes of sphere AuNPs (20 nm and 45 nm, respectively). The star peaked at 784 nm, and the rods’ LSPR peaks were in the near infrared (NIR) region at 660 and 687 nm.

**Figure 1.**
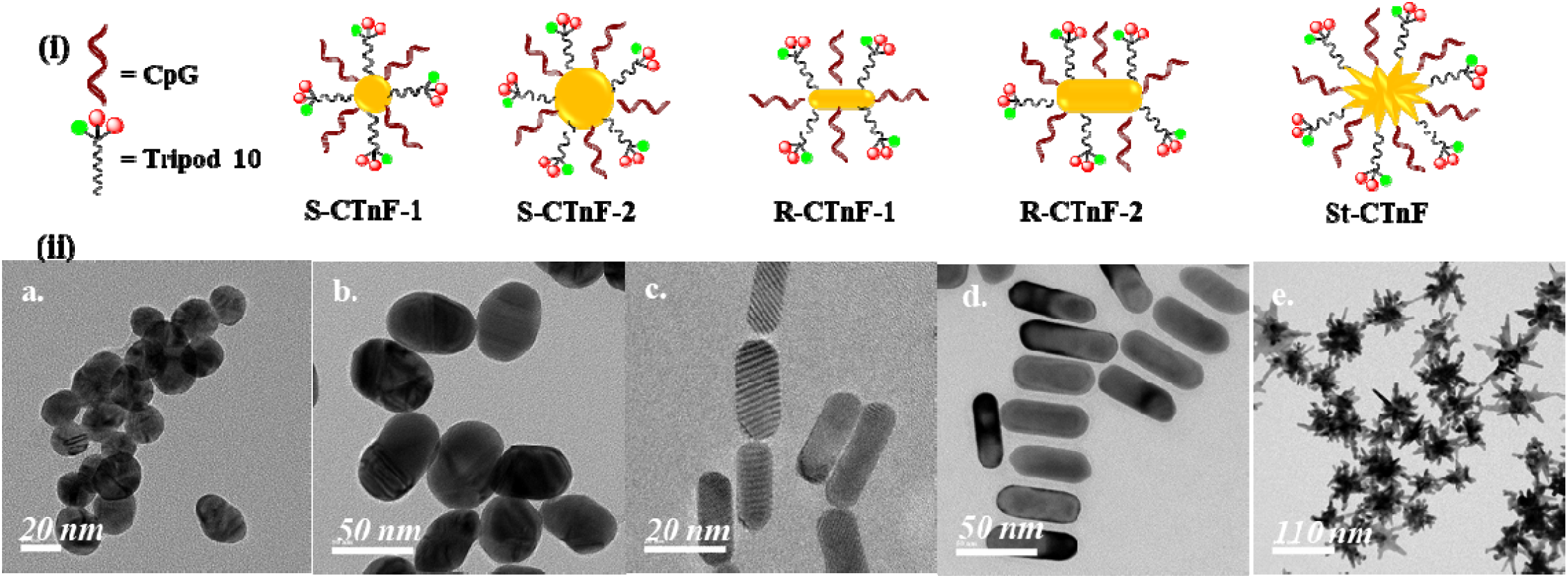
(i) Schematic representation of AuNPs conjugated with tripod **10** and CpG; (ii) TEM images of AuNPs **a**. S-CTnF-1, **b**. S-CTnF-2, **c**. R-CTnF-1, **d**. R-CTnF-2, **e**. St-CTnF

The AuNPs were encapsulated with the optimum number of CpG adjuvants, which is sufficient to induce immunomodulation by the ligand exchange process. Changes in the zeta (ζ) potentials of AuNPs confirmed the conjugation of the CpG adjuvant. More specifically, after CpG conjugation to rod AuNPs, the ζ-potential of rod AuNPs changed from a positive (+34.5 and +34.8, Table S1) to a negative potential (−11.4 and −17.7). In contrast, sphere and star AuNPs displayed slight changes in the negative potential, indicating effective CpG conjugation (Tables S2 and S3). Neither scanning electron microscopy (SEM) nor the UV-vis profile of these AuNPs showed any significant difference from the native nanostructures. The amount of CpG per AuNPs was quantified by a DNA quantification kit (Tables S2 and S3). Finally, tripod **10** was incorporated directly by mixing the CpG-AuNPs with the known quantity of ligand **10** to obtain **S-CTnF-1, S-CTnF-2, R-CTnF-1, R-CTnF-2**, and **St-CTnF**. The conjugation of tripod **10** was further confirmed by changes in the zeta potential and quantified using a thiol-detection kit. It was observed that, as the size and aspect ratio of the AuNPs increased, the number of CpG and tripod **10** per NP also increased because of the large surface area. As a control, we also synthesized and characterized five fluorescent CpG-adjuvant conjugated AuNPs (**S-CpG-1, S-CpG-2, R-CpG-1, R-CpG-2**, and **St-CpG**) and five TnF-antigen conjugated AuNPs (**S-TnF-1, S-TnF-2, R-TnF-1, R-TnF-2**, and **St-TnF**).

### Comparison of mDC uptake of nanostructures

To address the mechanism underlining the immunomodulation of the nanostructures, we first investigated the cellular internalization of AuNPs using mDCs because MGL (human macrophage galactose and N-acetylgalactosamine-specific C-type lectin) receptors interact with Tn-antigen on mDCs engaged receptor mediated endocytosis.^55^ Thus, it is hypothesized that disparity in the aspect ratio, nanostructure contact area, and antigen/adjuvant conjugation modulate the uptake mechanism and sequestration.

Since the nanostructures contain different concentrations of CpG and TnF conjugation, we adjusted the amount of AuNPs for *in vitro* and *in vivo* studies. We used 50 nmol of CpG as the optimum concentration for *in vitro* studies. Accordingly, the concentration of functionalized AuNPs was adjusted for experimental studies. Similarly, TnF concentration was quantified on CTnF-nanostructures and similar concentration of TnF-conjugated nanostructures were used for further biological studies. First, the cytotoxicity of AuNPs was investigated using an MTT assay employing mDC isolated from the spleen. As expected, none of the AuNPs showed cytotoxicity up to a concentration of 200 nmol of CpG (Figure S2). Then, AuNPs carrying 50 nmol of CpG concentration were incubated for 1 h and 4 h with mDC and imaged. The results indicated that rod-shaped AuNPs bearing TnF and CTnF ligands exhibited the most effective uptake compared with CpG conjugated AuNPs within 1 h. In contrast, sphere- and star-shaped AuNPs displayed weak uptake (Figure 2). This means that the shape of nanostructures promotes different rates of receptor-mediated endocytosis. We then quantified cellular uptake using FACS and developed a hierarchical clustering analysis (HCA). The HCA of the nanostructure displayed several distinct clusters. Although the Tn-antigen on the nanostructures increased the cellular internalization, the shape and size of the nanostructures displayed disparity in this cellular internalization; particularly, **R-TnF-1** showed strong uptake as compared to all other NPs (for 1 h, **S-TnF-1**: 50%; **S-TnF-2**: 30%; **R-TnF-1**: 80%; **R-TnF-2**: 30% and **St-TnF**: 60%), indicating rod-shaped nanostructures can pear the cell membrane *via* receptor mediated endocytosis and phagocytosis to induce strong uptake. Similar trend was also observed in CTnF-conjugated nanostructures. In contrast, sphere-shaped AuNPs showed weak and star shaped AuNPs showed moderate uptake. Among different size, **R-1** and **R-2** conjugates with CTnF and TnF showed a great disparity in the uptake rate, indicating that aspect ratio and ligand concentrations modulate the uptake rate. Furthermore, CpG-conjugated AuNPs resulted in the least uptake (Figure 2), probably due to formidable negative-negative charge repulsion between the cell-surface oligosaccharides and the oligonucleotides.^56^ This trend continued even at 4 h. Based on the results, we hypothesized that the origin of HCA is associated with the shape of the nanostructure and the interactions between the Tn antigen and C-type receptors on the cell surfaces. Moreover, These results correlate with previously published results with shape-dependent nanostructural cellular uptake.

**Figure 2.**
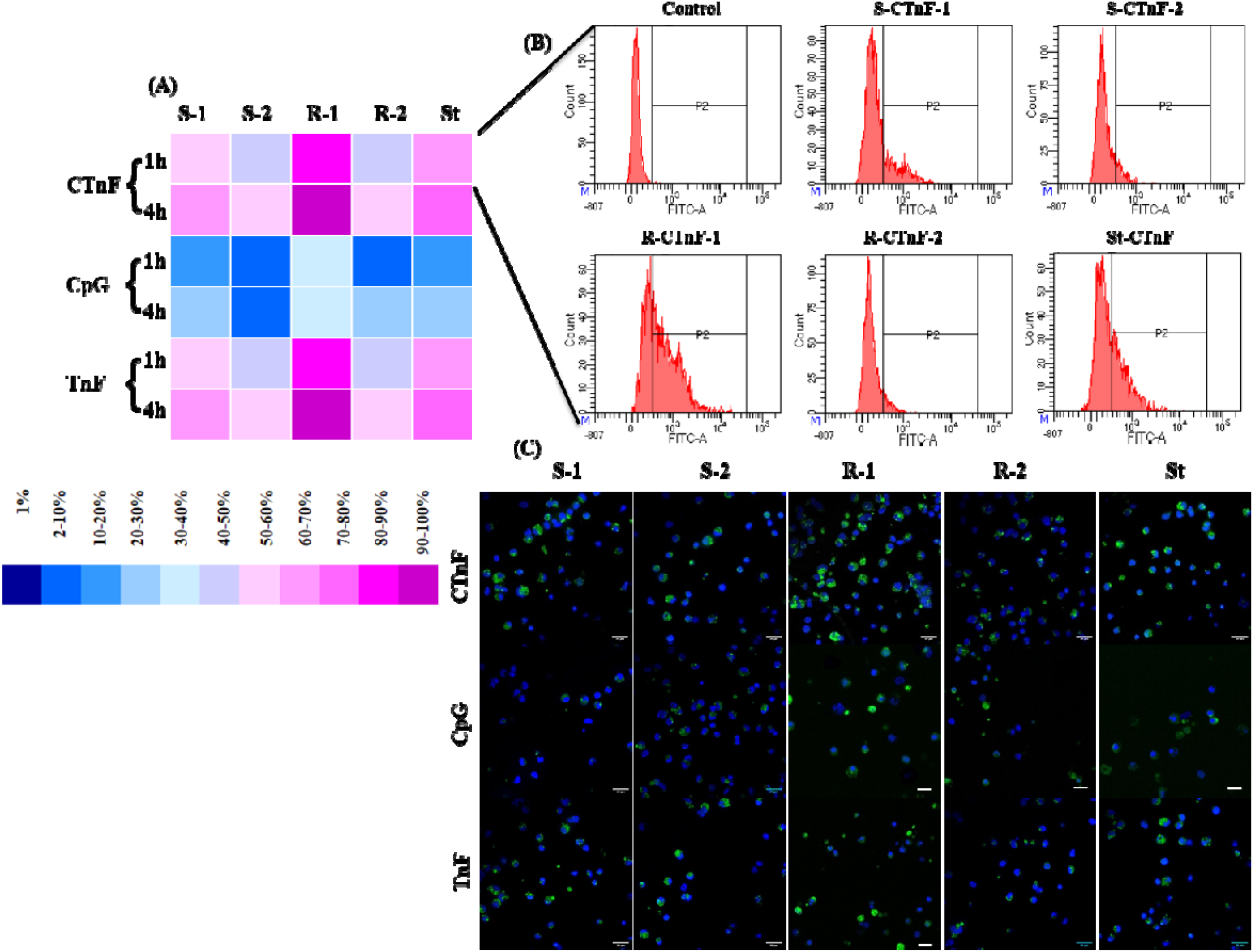
(**A**) Hierarchical clustering analysis (HCA) for cellular internalization of CTnF, TnF and CpG conjugated AuNPs by dendritic cells at 1 and 4 h interval based on FACS data (Uptake of R-CTnF-1 at 4h considered as 100%); (**B**) Flow cytometry data for CTnF conjugated AuNPs by dendritic cells after 1 h; (**C**) Confocal microscopy images for uptake of CTnF, TnF and CpG conjugated AuNPs by dendritic cells at 4 h (Scale bar 26 µm).

**Figure 3.**
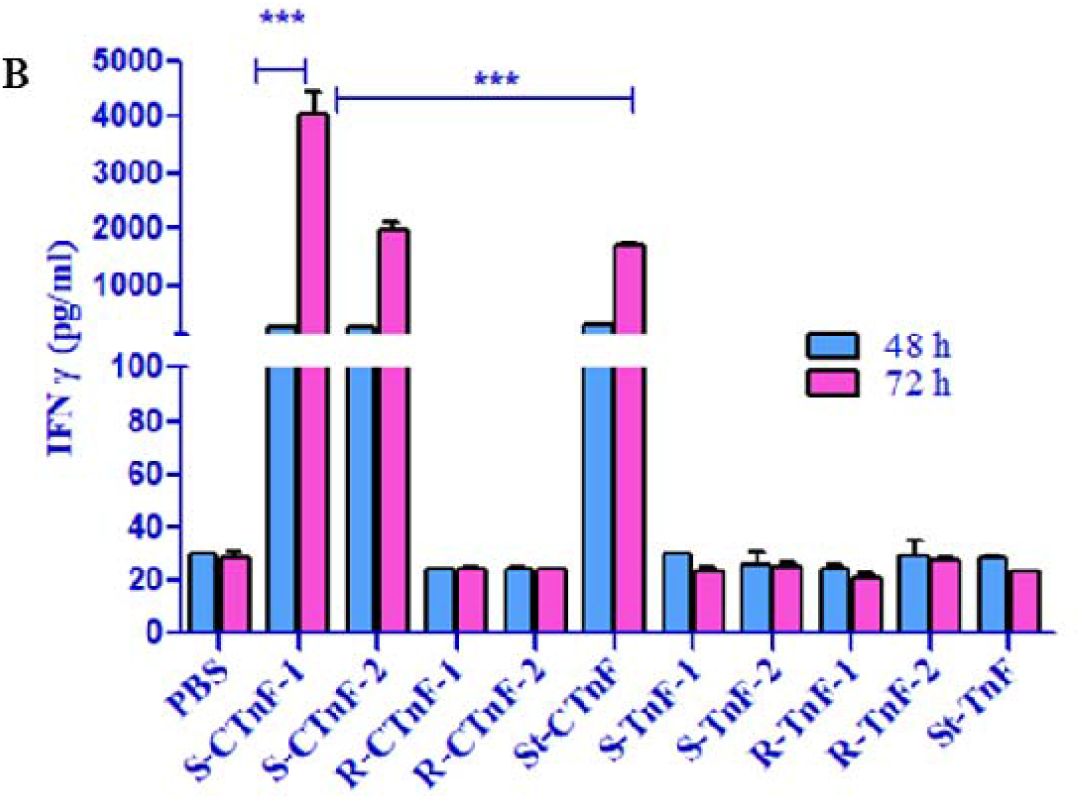
*In vitro* cytokine production by T cells in co-culture with dendritic cells pulsed with CTnF and TnF conjugated AuNPs. Interferon-γ (IFN-γ) production after 48 and 72 h; Asterisks indicate statistically significant differences (\*\*\**p* <0.0001)

**Figure 4.**
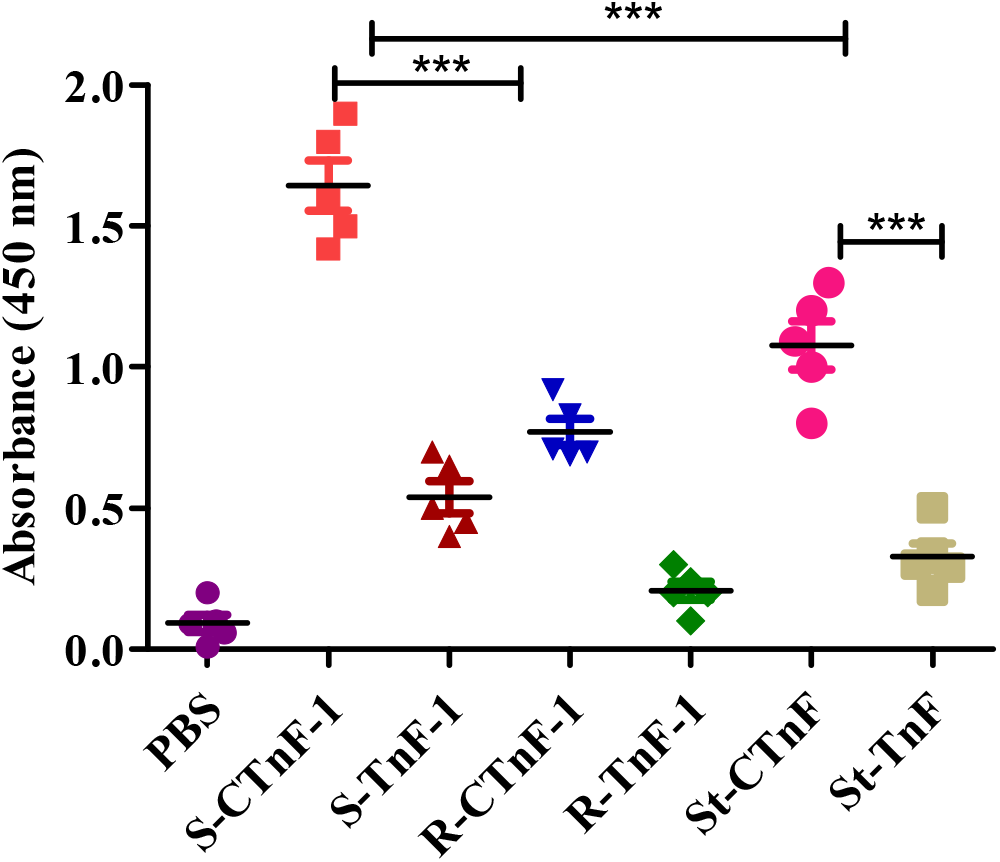
Total IgG antibody titer after immunization of C57BL/6 mice with CTnF and TnF functionalized sphere and star AuNPs. Asterisks indicate statistically significant differences (\*\*\**p* <0.000)

### Inflammatory activation in DC/T-cell co-culture assay

Upon cellular internalization of the nanostructure, the CpG adjuvant activates the DCs and processes the antigens as a major histocompatibility complex (MHC), resulting in the activation and differentiation of T-cells into Th1, Th2, and Th17 cells. To analyze these adaptive immune responses, we performed a DC/T-cell co-cultivation assay and quantified the level of cytokines secreted in the media in the presence of nanostructures.^57^ Stimulation of mDC/T-cells with the nanostructure resulted in nanostructures’ shape and adjuvant-dependent enhanced secretion of interleukin (IL)-6, interferon (IFN)-γ, and tumor necrosis factor (TNF)-α; however, IL-10 (Th2-type immune response) was poorly secreted. The cytokine secretion analysis of the AuNPs indicated that CTnF-conjugated AuNPs displayed a stronger Th1-cytokine response than the TnF-conjugated AuNPs, confirming that antigen/adjuvant conjugated synergistically modulates T-cell activation.

Interestingly, the rod-shaped AuNPs showed weak Th1-cytokine secretion as compared to sphere and star shaped AuNPs. This outcome is not entirely surprising, since this type of disparity in cytokine secretion has been reported for nanostructures. Previously, Niikura *et al*. reported the effect of nanostructure shapes on displaying different rates of sequestration in lysosomes and cytosols to modulate immune responses, and ultimately, antibody production.^28^ Lee *et al*. compared sphere and star-shaped AuNPs sequestration in the endosomes of macrophages and showed that the disparity in the assembly correlates with immune activation.^58^ In our case, the **R-CTnF-1** impeded TLR9 receptor activation even though they exhibit a great degree of cellular internalization. This may be due to the inherent self-assembly nature of rod-shaped AuNPs and the different rates of sequestration of AuNPs in lysosomes, cytosol, and the endosomal region (Figure S4). Together, these results reiterate that the cellular uptake mechanism and immunomodulation are two independent mechanisms, and the shape of the nanostructures is a crucial factor in controlling the immune process.

### Antibody responses against synthetic Tn-peptide

As a final component of the study, the *in vivo* immunogenicity of nanostructures was examined in the C57BL/6 mice model. A groups of five mice were subcutaneously immunized with 100 µl of TnF and CTnF functionalized sphere-, rod- and star-shaped nanostructures (containing 5 nmol of CpG and 7–10 μg of Comp **10**) on days 0, 14, and 28. On day 36, the mice were sacrificed; their serum was harvested, and the IgG titer specific to compound **6** was determined by enzyme-linked immunosorbent assay (ELISA). Significantly, **S-CTnF-1, R-CTnF-1** and **St-CTnF** showed the higher antibody induction compared with **S-TnF-1, R-TnF-1** or **St-TnF** nanostructures. Notably, **S-CTnF-1** showed the highest immune responses of all six structures. Among three **CTnF** conjugated AuNPs, **R-CTnF-1** showed least IgG titer. These results indicate that the shape of AuNPs has a pivotal role in modulating immune responses and further vaccine development.

## Conclusion

Different shapes and sizes of AuNPs were used to fine-tune the immune responses of the Tn antigen in the presence of CpG adjuvant. Validated by a series of imaging techniques, cytokine secretion and *in vivo* antibody secretion studies, the results showed that cellular internationalization and cytokine secretion are two independent mechanisms. Furthermore, although sphere- and star-shaped nanostructures showed ineffective cellular uptake compared with rod-shaped nanostructures, they were observed to secrete Th1 bias immune responses, resulting in a potential platform to develop TD immune modulation of weak immunogenic antigens. Finally, the simplicity and effectiveness of nanotechnology may hold the key to accelerating glyconanotechnology in vaccine research.

## Experimental Methods

### Confocal Imaging

Dendritic cells (2 × 10^6^ cells) were seeded on poly-D-lysine coated coverslip in complete IMDM medium and incubated at 37 °C for overnight. The cells were incubated with CTnF, CpG and TnF functionalised AuNPs (50 nmol of CpG) for different time intervals 1 h and 4h. Then the cells were washed with the cold PBS and fixed with 4% paraformaldehyde at room temperature for 20 min. Next, the coverslips were washed with PBS and water and mounted on a slide using medium (Vectashield). The fluorescent images were taken using Lieca sp8 microscope.

### FACS Analysis

Dendritic cells (2 × 10^6^ cells) were seeded in 96 well plates in IMDM media and incubated for 30 min. The cells were pulsed with CTnF, CpG and TnF functionalized AuNPs (50 nmole of CpG) for 1 h and 4h. Then the cells were washed with cold PBS and resuspended in FACS buffer and proceeded for analysis. Quantification of uptake was done by using Flowjo software.

### mDC/T-cell Co-culture assay

Dendritic cells (2×10^6^ cells/well) were seeded in 96 well-plate in IMDM medium at 37 °C for 30 min. Then cells were pulsed with the CTnF and TnF functionalised AuNPs (50 nmol of CpG) and incubated further for an hour. Next, the purified T-cells (60 µl of 7×10^6^/well) were added and incubated for 48 and 72 h. Cytokine (IL6, IL10, TNFα, IFNγ) level in the supernatant was determined after 48 and 72 h of stimulation using ELISA (R&D Systems).

### Immunization protocol

Immunization studies were carried out using 6 to 8 weeks old females (C57BL/6) mice as per protocol approved by the institutional ethical committee. Five groups of mice (n=5) were immunized subcutaneously with 100 µl of each S-CTnF-1, R-CTnF-1, St-CTnF, S-TnF-1, R-TnF, St-TnF (contains 7 to 10 µg of Tn and 5 nmol of CpG) and PBS on days 0, 14, 28. The mice were bled on days 36 and serum antibody titer was analysed by ELISA.

### ELISA for evaluation of IgG antibody titer

The IgG antibody titers were quantified using maleic anhydride activated 96 well plates.^59^ The plates were coated with Tn glycopeptide (Comp **6**) (3 µg/ml in 0.1 M carbonate buffer pH 9.2) at room temperature for 1 h followed by incubation at 4 °C for overnight. The coated plates were washed with washing buffer (PBS + 0.1 % Tween-20) and blocked with blocking buffer (2 % BSA) for 1h at room temperature. Next, the plates were washed, mice anti-serum (1:1000 diluted) was added and incubated for 2 h at room temperature followed by washing. HRP-coated secondary goat anti-mouse antibody (1:2000 diluted, Thermofisher) was added to the plate and incubated for 2 h at room temperature. The plates were washed, TMB substrate solution was added, and the reaction was stopped by adding 0.5 M H_2_SO_4_. The absorbance was recorded at 450 nm.

## ASSOCIATED CONTENT

### Supporting information

Information about used materials and synthesis, as well as detailed analytical data of the compounds and data from the biophysical are available. The Forting Information is available free of charge on the ACS Publications website at

## Author Contributions

All authors have given approval to the final version of the manuscript.

## ACKNOWLEDGMENT

R. K. gratefully acknowledges financial support from the IISER, Pune, DBT (grant nos. BT/PR21934/NNT/28/1242/2017, BT/PR34475/MED/15/210/2020), STARS/APR2019/CS/426/FS and SERB/F/9228/2019–2020. P.M.C. acknowledge DBT-*BioCARe* grant BT/PR19727/101/253/2016 for financial support and S.T. acknowledges CSIR-SRF, EMBO Short-Term Fellowship.

---

**Scheme 1.**
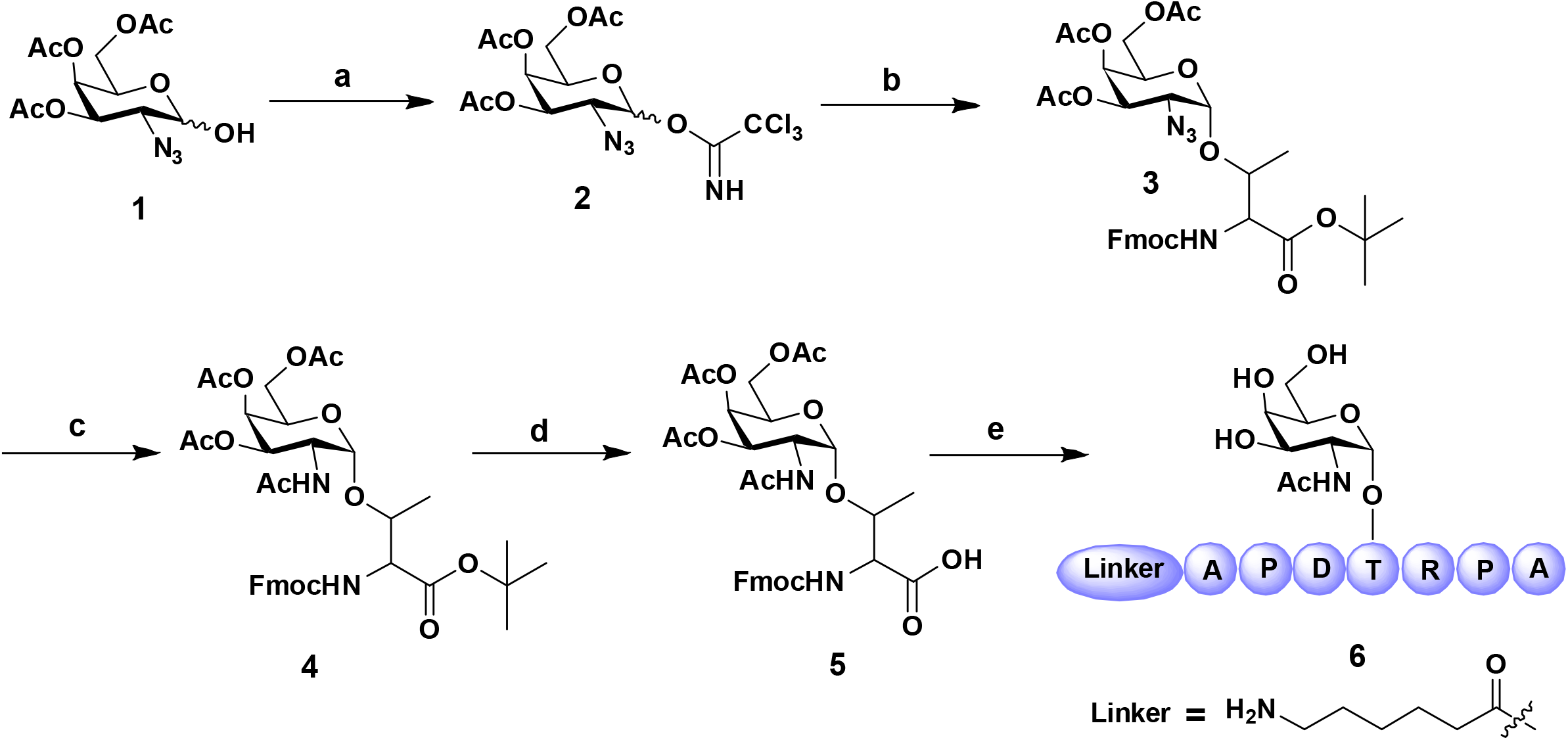
Reagents and conditions: (a) DBU, Cl_3_CN, CH_2_Cl_2_, rt, 2 h, 88%; (b) TMSOTf, Fmoc-Thr-OtBu, CH_2_Cl_2_, rt, 1 h, 71%; (c) Zn, THF/AcOH/Ac_2_O (3:2:1, v/v), rt, 4 h, 61%; (d) TFA/DCM (1:1, v/v), rt, 4 h; (e) Solid phase peptide synthesis and complete deprotection.

**Scheme 2.**
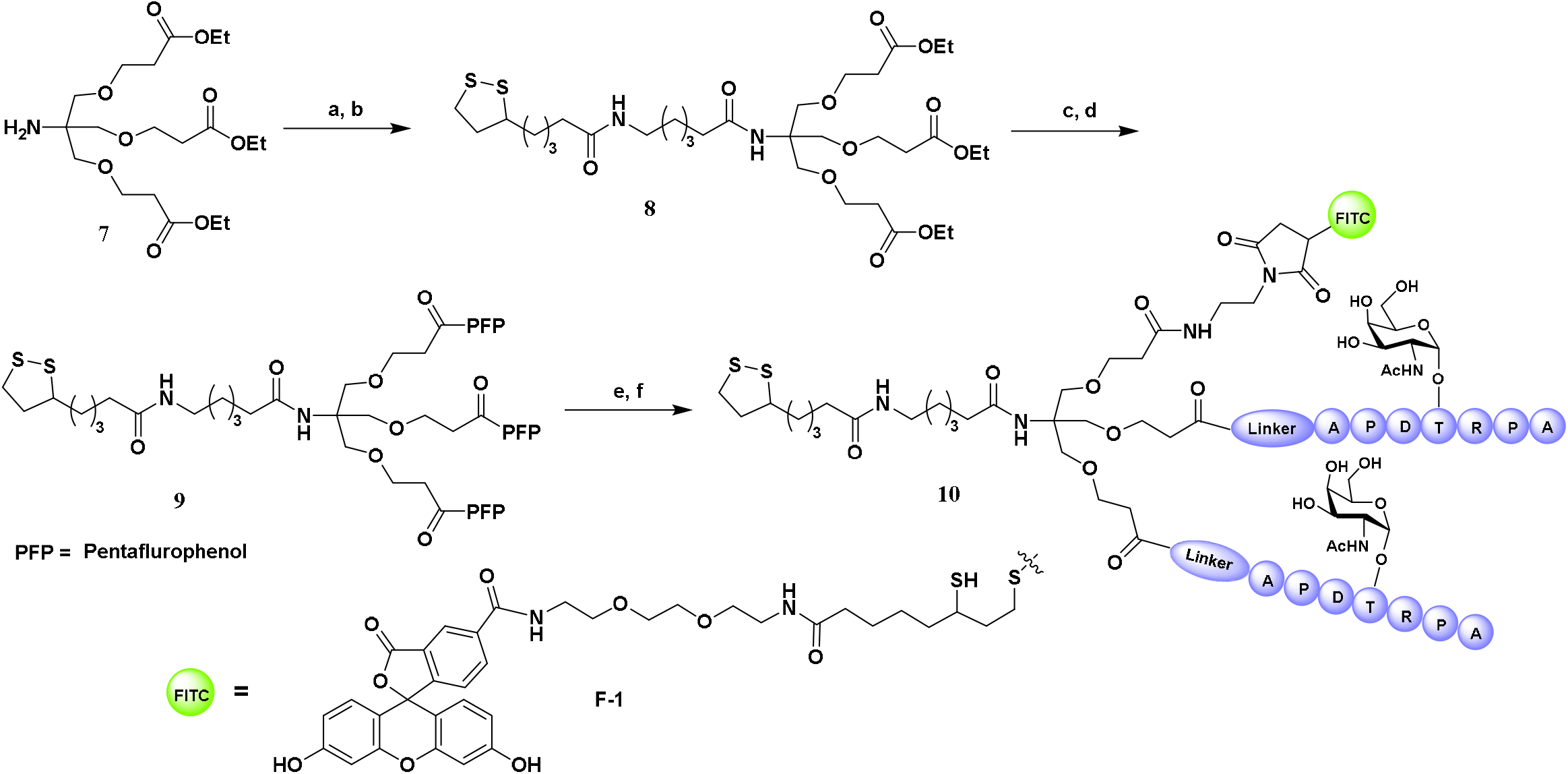
Reagents and conditions: (a) DCC, 5-(Boc-amino)pentanoic acid, CH_2_Cl_2_, rt, 12 h, 69%; (b) i) CH_2_Cl_2_/TFA (3:1, v/v), rt, 4 h; ii) DIPEA, N-Lipoyloxy succinimide, CH_2_Cl_2_, rt, 4 h, 82% over two steps; (c) i) LiOH, THF/H_2_O/Dioxane (2:1:1, v/v), rt, 6 h; d) Pentaflurophenol, DCC, CH_2_Cl_2_, rt, 12 h, 47% over two steps; (e) i) *N*-(2-Aminoethyl)maleimide trifluoroacetate, DIPEA, CH_2_Cl_2_, rt, 2 h, 31% ; ii) Tn glycopeptide, DIPEA, DMF, rt, 4 h, 58%; (f) FITC linker, DIPEA, DMF, rt, 24 h, 16%.

